# Handling Noise in Protein Interaction Networks

**DOI:** 10.1101/527606

**Authors:** Fernanda B. Correia, Edgar D. Coelho, José L. Oliveira, Joel P. Arrais

## Abstract

Protein-protein interactions (PPI) can be conveniently represented as networks, allowing the use of graph theory in their study. Network topology studies may reveal patterns associated to specific organisms. Here we propose a new methodology to denoise PPI networks and predict missing links solely based on the network topology, the Organization Measurement (OM) method. The OM methodology was applied in the denoising of the PPI networks of two *Saccharomyces Cerevisiae* datasets (Yeast and CS2007) and one *Homo Sapiens* dataset (Human). To evaluate the denoising capabilities of OM methodology, two strategies were applied. The first compared its application in random networks and in the reference set networks, while the second perturbed the networks with the gradual random addition and removal of edges. The application of OM methodology to the Yeast and Human reference sets achieved an AUC of 0.95 and 0.87, in Yeast and Human networks, respectively. The random removal of 80% of the Yeast and Human reference sets interactions resulted in an AUC of 0.71 and 0.62, whereas the random addition of 80% interactions resulted in an AUC of 0.75 and 0.72, respectively. Applying the OM methodology to the CS2007 dataset yields an AUC of 0.99. We also perturbed the network of the CS2007 dataset by randomly inserting and removing edges in the same proportions previously described. The false positives identified and removed from the network varied from 97%, when inserting 20% more edges, to 89% when 80% more edges were inserted. The true positives identified and inserted in the network varied from 95% when removing 20% of the edges, to 40% after the random deletion 80% edges. The OM methodology is sensitive to the topological structure of the biological networks. The obtained results suggest that the present approach can efficiently be used to denoise PPI networks.

## Introduction

Proteins are central players in every organism, as they are required for virtually every single cellular function. However, proteins are required to interact with one another to fulfil their functions. For this reason, disease states may appear, if the physiological interaction between two proteins is disrupted [1].

Protein-protein interactions (PPI) networks are a subset of complex biological networks that have specific topological properties, such as a high clustering coefficient, the presence of hierarchy, heterogeneity and a power-law-like degree distribution [2]. The guilt-by-association hypothesis states that two proteins sharing many interactive neighbours are likely to hold functional homogeneity and localization coherence [3]. These characteristics suggest that network topology alone may be a viable option for PPI network denoising. PPI network denoising corresponds to find interactions that do not exist and to find missing interactions. Methods to determine protein interactions are not accurate and organisms are not yet fully known, being important to denoise PPI networks to have more precise models of the organisms.

Protein interactions can be represented as graphs, allowing the use of graph theory in their study. As such, different methods were developed to denoise biological networks [reviewed in 4]. These include repeating experiments [5, 6], using prior knowledge about proteins [7, 8], using functional or structural annotations [9-13] and using comparisons with theoretical distributions constructed from known data and network topology-based approaches [14-18]. The herein proposed approach falls under the latter category.

An approach, called Non-Convex Semantic Embedding (NCSE) evaluates the reliability of interactions in a PPI network trying to learn a Euclidean embedding under the geometric assumption of PPI networks [19] and it was tested in three data sets of *Yeast Saccharomyces Cerevisiae*. Computational methods based on machine learning were also applied to evaluate the reliability of PPI and tested in *Yeast* and *Helicobacter Pylori* PPI datasets [20, 21].

In a recent study, Lü et al. [22] proposed the structural consistency index and the Structural Perturbation Method (SPM). On the one hand, the structural consistency index can reflect the inherent link predictability of a network without knowing its organization a priori, allowing to estimate the explicability of the organization of a network, and to supervise mechanistic changes during the evolution of the network. On the other hand, the SPM performs link prediction by removing a percentage of the edges in a network, thus perturbing the remaining network by that percentage. This is based on the strong correlation between independent network perturbations, which suggests that the missing edges, i.e., False Negative (FN) interactions can be identified by perturbing the networks with an additional set of known interactions, i.e., True Positive (TP) interactions.

Luo et al. [4] proposed the Collaborative Filtering-enhanced Topology-based (CFT) method to perform protein interatomic mapping on sparse High-Throughput Screening (HTS) PPI data, since the performance of network topology-based approach usually deteriorates when using sparse network data. This approach is based on the notion that the solution space of the interatomic mapping and the solution space of the personalized recommendations are similar. Each protein is represented as a feature vector that describes their interactions in the network. In addition, the feature vector is used to calculate the corresponding similarity vector that represents the interactions through the Functional Similarity weight, creating an Inter-Neighbourhood Similarity (I-Sim) for modelling PPIs. Functional parameters for each protein in the dataset are obtained from Gene Ontology (GO), allowing the use of functional similarity measures. Denoising of the input HTS-PPI data is performed via the integration of saturation-based strategies into the I-Sim, achieving a precise relationship model. Their method was applied to three different datasets and compared with three other algorithms (Interaction Generality [14], Czekanowski-Dice distance [15], and Functional Similarity weight [16]), showing better performance on large, sparse HTS-PPI datasets. Since they use GO annotations to characterize their proteins, this approach is likely to underperform when considering less studied organisms.

A different strategy termed Intrinsic Geometry Structure (IGS) was proposed by Yi Fang et al. [23]. IGS is a geometry-based approach which uses heat diffusion in the PPI network to collect structural information about all paths connecting two given nodes, thus defining intrinsic relationships among them. They use a maximum likelihood-based algorithm to determine the optimal dissipation time, predicting the global structure of the PPI network from the local structure. After performing heat diffusion for the optimal dissipation time, the intrinsic geometric structure of the PPI network is revealed. One of the main advantages of the IGS method is its robustness against missing protein associations and sparse PPI data. Their method was tested with the *S. cerevisiae* (CS2007) network [24], a network of the bottlenose dolphin community [25], and a network of known terrorist cells [26]. In addition, they compared the performance of IGS with two other methods, the Multi-Dimensional Scaling-based (MDS) method [27] and the Hierarchical Random Graph (HRG) method [28], showing that IGS performed slightly better than MDS when tested with the CS2007 dataset and better than IGS and HRG for the other datasets tested. Their analysis was based only in the Area Under the Receiver Operating Characteristic (ROC) Curve (AUC) values.

Among the described works, the MDS method proposed by Kuchaiev et al. [27] is the only one relying in PPI network topology that was applied in a Homo Sapiens PPI dataset. To address the sparsity problem of the networks, Luo et al. [4] uses collaborative filtering, but the method was not tested on perturbed networks, i.e., when added random noise. The IGS method [23] was compared to the MDS method using the Yeast Saccharomyces Cerevisiae PPI dataset, and both methods were tested in perturbed networks with the same percentages (from 10% till 80%). However, they were not able to determine which of the edges recovered belong to removed group, or which of the edges removed are then recovered.

In this paper, we introduce the Organization Measurement (OM) method to denoise PPI networks based exclusively in the network topology. Topological measures are used to find trends that characterize interacting and non-interacting proteins distributions. A high confidence set of protein interactions is used to construct a network, followed by the calculation of the weights of interactions and non-interactions in the network. The OM weighted matrix is obtained and used to find distribution trends that allow to distinguish interaction distributions from non-interaction distributions. The OM threshold value that better distinguishes these types of distributions is then used to identify False Positive (FP) interactions and FN (novel) interactions. This way, an OM topological model is built to be used in the denoising of a network, resulting in a better approximation of the expected network.

## Materials and Methods

This section will describe how we obtained the datasets used in the experiments and the topological measures used with the Organization Measurement (OM) methodology, including the new Neighbourhood Clustering (NC) proposed measure. It will also describe the OM methodology, how to obtain the OM matrix of weights and how to determine the threshold value, giving a description of the OM methodology pipeline (Fig 1) to denoise networks.

**Figure 1:**
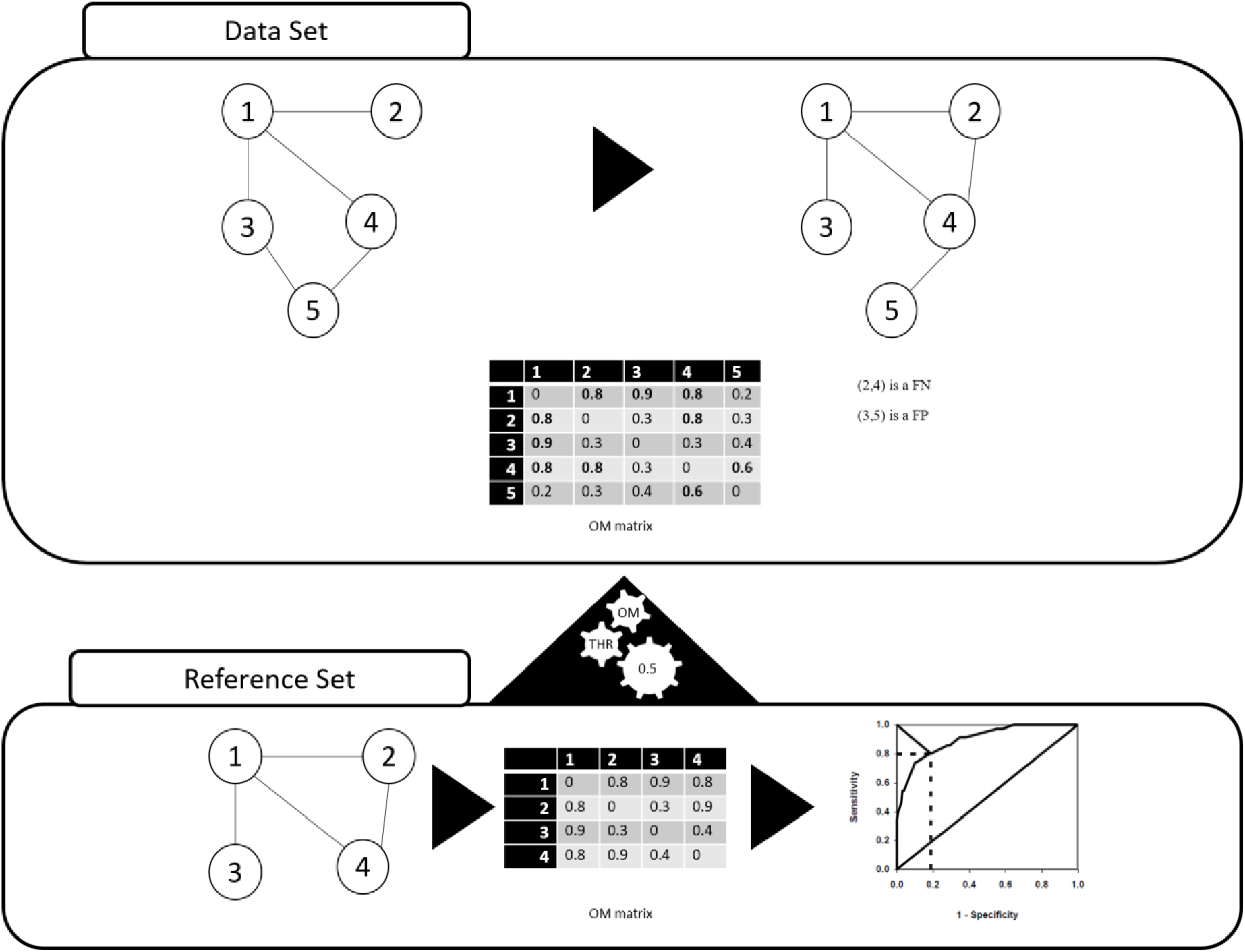
Diagram of the OM methodology pipeline. A Reference Set of an organism is used to create a network model and the respective OM matrix is obtained, using the topological properties. The threshold (OM THR) is calculated using the ROC curve and choosing the value that best separates interactions from non-interactions distributions. OM THR is then applied to denoise a lower-confidence Data Set network of the same organism, using the respective OM matrix.

For each organism, we collected a set of high-confidence PPI interactions. Although these PPI do not reflect the entirety of the protein interaction networks of the selected organisms, they were used to construct the known PPI network of each organism, *i.e.*, their reference sets.

In the application of the OM methodology, various topological measures were calculated to characterize these networks, based on the assumption that these measures will allow the identification of topologic patterns to support network denoising. In this paper, we use the term “denoising” to define the identification of FP and FN interactions, removing the former cases from the network while adding the latter. The methodology proposed here can also be used to rank the level of confidence of the interactions already presented in the network. Different topological measures can identify different patterns and thus, here we consider that different topological measures can contribute to the denoising process.

Next sections will describe in a detailed way the OM methodology pipeline, after the description of the datasets chosen to illustrate and validate the proposed methodology and the description of the topological measures used to test OM methodology, including the new one topological measure proposed. A section will explain the process to obtain the weighted OM matrix from the adjacency matrix, another section will describe how to determine the OM THR value using the ROC curve and a last section will describe how to use the threshold in the denoising of PPI networks.

### Datasets

The Search Tool for the Retrieval of Interacting Genes/Proteins (STRING) database [29] contains known and predicted protein interactions of various organisms. PPI in STRING derive from five main sources: (1) genomic context predictions; (2) high-throughput experimental methods; (3) conserved co-expression experiments; (4) automated text mining, and; (5) previous knowledge from third-party databases. Each interaction in STRING has an associated score for each prediction method and has a Combined Score (CS) that range from 0 to 1000, indicating the degree of confidence of each interaction. Calculation of the CS considers several parameters, such as the number and the quality of different sources indicating that a PPI occurs.

The interactions derived by experimental methods with a score greater than 900 have been considered of high-confidence in multiple works [30, 31]. Therefore, the reference sets used in this work comprise experimentally determined PPI data obtained from STRING with a score greater than or equal to 900.

These data were collected from two different organisms, namely the *Yeast Saccharomyces Cerevisiae* (Yeast) and *Homo Sapiens* (Human) [29]. Using these data, an undirected network is constructed for each organism and the main component is extracted.

Table 1 summarizes the characteristics of the reference set networks obtained for Yeast and Human, including the number of nodes, the number of edges, the average degree and network density. The observed average degree and density values are highly suggestive that these are biological networks are sparse, *i.e.*, they have much less edges than the complete network with the same set of nodes. Our high-confidence networks (*i.e.*, PPIs obtained from the STRING database with experimental source score greater than 900) comprised 29,319 interactions between 3,937 proteins for the Yeast dataset and 16,931 interactions between 4,943 proteins for the Human dataset.

**Table 1.** Topological characteristics of the Yeast, CS2007 and Human networks used as reference sets.

| Organism | Name | From | N° of nodes | N° of edges | Average degree | Density |
| --- | --- | --- | --- | --- | --- | --- |
| Yeast<br>*Saccharomyces Cerevisiae* | Yeast | STRING experimental<br>CS >= 900 | 3,937 | 29,319 | 14.8941 | 0.0038 |
|  | CS2007 | compiled by<br>Collins *et al.* | 1,004 | 8,323 | 16.5797 | 0.0165 |
| Homo Sapiens | Human | STRING experimental<br>CS >= 900 | 4,943 | 16,931 | 6.8505 | 0.0014 |

Additionally, we used a high confidence external dataset compiled by Collins *et al.* [24] and referred to as CS2007 hereafter, to compare the proposed methodology with other topology-based denoising methods [23, 27, 32]. This dataset comprises 9,074 PPIs between 1,622 unique proteins from *Yeast Saccharomyces Cerevisiae*. To ensure a direct comparison between the OM methodology and the existing methods we followed their approaches and only used the largest connected component. The largest connected component of the dataset compiled by Collins *et al.* [24] includes 8,323 interactions between 1,004 proteins (Table 1).

### Measures for similarity and diversity analysis of network data

Protein interactions can be conveniently modelled as a network, where each node represents a protein and each edge represents an association between two proteins. The most commonly used technique to quantify the interaction profile similarity of protein interaction networks (or any type of biological network) relies on association indices. Bass *et al*. [33] performed a comprehensive review on the selection of association indices for the analysis of gene similarity. In their work, the Jaccard (JC), Geometric and Cosine indices were shown to be the most versatile, as though not excelling in any particular task, their strengths were the most balanced out of all evaluated measures. A review of similarity indices can also be found in [34]. Daminelli *et al*. test the application of different association indexes to bipartite networks [35].

A more recent study reports that the JC measure performs better than three other measures in a specific model [36].

The JC measure is defined as the ratio of the intersection of the number of neighbours of nodes *i* and *j* divided by their union (*i.e.*, the ratio of nodes shared between *i* and *j* divided by the total number of nodes connected to both):

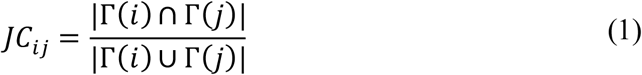

where Γ(*i*) is the set of neighbours of *i.* We also explored and tested additional measures and two of them that gave good results were Betweenness (BETW), and Katz indexes.

The implementation of Betweeness (BETW) used was:

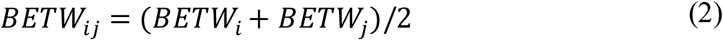

where

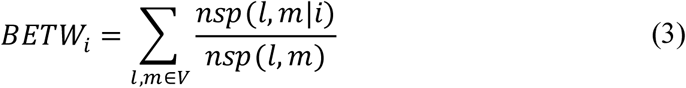

where *V* is the set of nodes, *nsp*(*l, m*) is the number of (*l, m*) shortest paths and *nsp*(*l, m*|*i*) is the number of those paths passing through the node *i*.

The implementation of Katz used was:

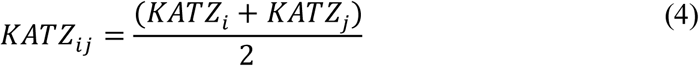

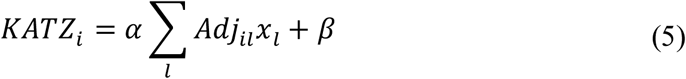

with *Adj* the adjacency matrix of the network with eigenvalues λ. It was used *α* = 1/λ_*max*_ and *β* = 0, when Katz centrality is the same as the eigenvector centrality.

Based on the idea that closely associated proteins are more likely to interact, that the network modularity is associated with the Clustering Coefficient (CC) [37] and a high mean CC of a community can be used to identify those that are functionally homogeneous [38], we implemented a novel measure to emphasize the relevance of the CC concept associated to the neighbourhood concept in a network. This measure was called Neighbourhood Clustering (NC) measure and is defined as the ratio of the sum of the CC of the nodes shared between *i* and *j* divided by the sum of the CC of the total number of nodes connected to both *i* and *j*:

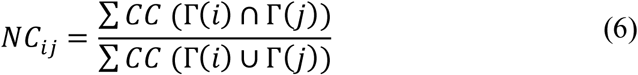

where Γ(*i*) is the set of neighbours of *i*.

### Organization measurement matrix

In Fig 1 we summarize the pipeline of the proposed OM methodology. Once the PPI network (Reference Set) for the or’ganism is constructed, its respective adjacency matrix is built, followed by its transformation into a weighed matrix, the OM matrix. The OM matrix is used to find distribution trends that allow to distinguish interactions and non-interactions. The weights for interactions and non-interactions are calculated using topological measures and using the information about the interactions of the network.

The adjacency matrix of the PPI network A, with *N* proteins and *M* interactions is defined as *adj*_*A*_ = [*a*(*i, j*)], where *a*(*i, j*) = 1, if there is an interaction in A between nodes *i* and *j*. Otherwise, *a*(*i, j*) = 0. A topological measure is applied to A to determine a weight for each (*i, j*) to transform the adjacency matrix *A* into a transformed matrix *A*_*w*_ = [*a*_*w*_(*i, j*)], where *a*_*w*_(*i, j*) is the weight of (*i, j*) in *A*, calculated using the topological properties of the network.

The weight *a*_*w*_ (*i, j*) represents the strength value of the edge (*i, j*) per the topological measure used and aims to capture patterns associated to the network that can originate signatures that identify the PPI network of each organism. This weight was used to characterize interaction and non-interaction distributions of the PPI network to determine the separation border between them.

### Organization measurement threshold value determination

One of the assumptions made in this work is that the PPIs in the reference datasets are true. This assumption can be made due to the sparsity of protein interaction networks and the rigorous criteria chosen to filter TP interactions. However, the same cannot be said for the non-interactions, as the presence of FN PPI is highly likely.

The value that best distinguishes both interactions and non-interactions distributions was called the OM threshold value. First, we collect protein interactions data of a specific organism and then a network is built (Fig 1). Then, the respective adjacency matrix is constructed, followed by its transformation into a weighted matrix, the OM matrix, using the topological measures of interest. Finally, the ROC curve is calculated and used to determine the optimal cut-off, corresponding to the threshold value that separates the interaction distributions from non-interaction distributions. We considered as the optimal cut, the point closest to (0,1) in the ROC curve, where sensitivity equals specificity. Different topological measures were tested and the respective cut-off values were determined. The outcomes of these experiments are described in the “Results and discussion” section.

### Organization measurement methodology to find spurious and new interactions

An accepted assumption in network topology-based approaches is that interacting proteins in a local community and closer to one another in the network are most likely involved in similar functions, or part of the same pathways [39-41]. The use of topological measures that capture this information should be prioritized, as they are expected to better grasp patterns in incomplete networks, thus allowing the approximation of incomplete input networks to the real networks.

From each reference set (Reference Set) PPI network we calculated its adjacency matrix. Then, after calculating the respective weights, the adjacency matrix is transformed into a weight matrix. Finally, the threshold that best separates PPI and non-PPIs was determined through the finding of the optimal cut-off of the ROC curve. This threshold was applied to detect spurious and missing PPI in the network (Data Set), to obtain a better approximation of the true network. In the example network shown in Fig 1, there are five nodes representing five different proteins, in addition to six edges that could represent the interactions between them (Data Set). Assuming the example network approximates the current knowledge of a given biological network, not all true interactions are represented and the existence of FP is expected. Once the threshold value is calculated, using the reference set (Reference Set), it is applied to the OM matrix of the Data Set, to identify FP and FN interactions. FP interactions are then removed from the network, whereas FN interactions are added.

## Results and Discussion

The OM methodology was tested with different topological measures and was evaluated using three different scenarios. Next subsections will describe the results obtained with the experiments made.

### Analysis of different topological measures to identify the optimal threshold value

A key component of the proposed method is the determination of the threshold value to discriminate between protein interactions and non-protein interactions distributions. As such, we decided to test the OM methodology with different topological measures to determine which better discriminates PPI from non-PPI. The four topological measures used were the JC, the BETW, the KATZ and the proposed new measure, the NC. Fig 2 shows the ROC curves when using this methodology with four different topological measures for the Yeast and Human organisms and Fig 3 shows the same information, but after data normalization between 0 and 1. Table 2 shows the respective AUC values obtained. Best results were achieved when using OM methodology with the JC and NC measures.

**Table 2:** AUC in Yeast and Human datasets.

| Table 2: AUC in Yeast and Human datasets. |  |  |  |  |
| --- | --- | --- | --- | --- |
| Topological | Yeast AUC |  | Human AUC |  |
| measures | Not normalized | Normalized | Not normalized | Normalized |
| JC | 0.9462 | 0.9460 | 0.8438 | 0.8458 |
| BETW | 0.7142 | 0.7676 | 0.7659 | 0.8004 |
| KATZ | 0.7151 | 0.7714 | 0.7258 | 0.7944 |
| NC | 0.9534 | 0.9526 | 0.8708 | 0.8700 |

**Figure 2.**
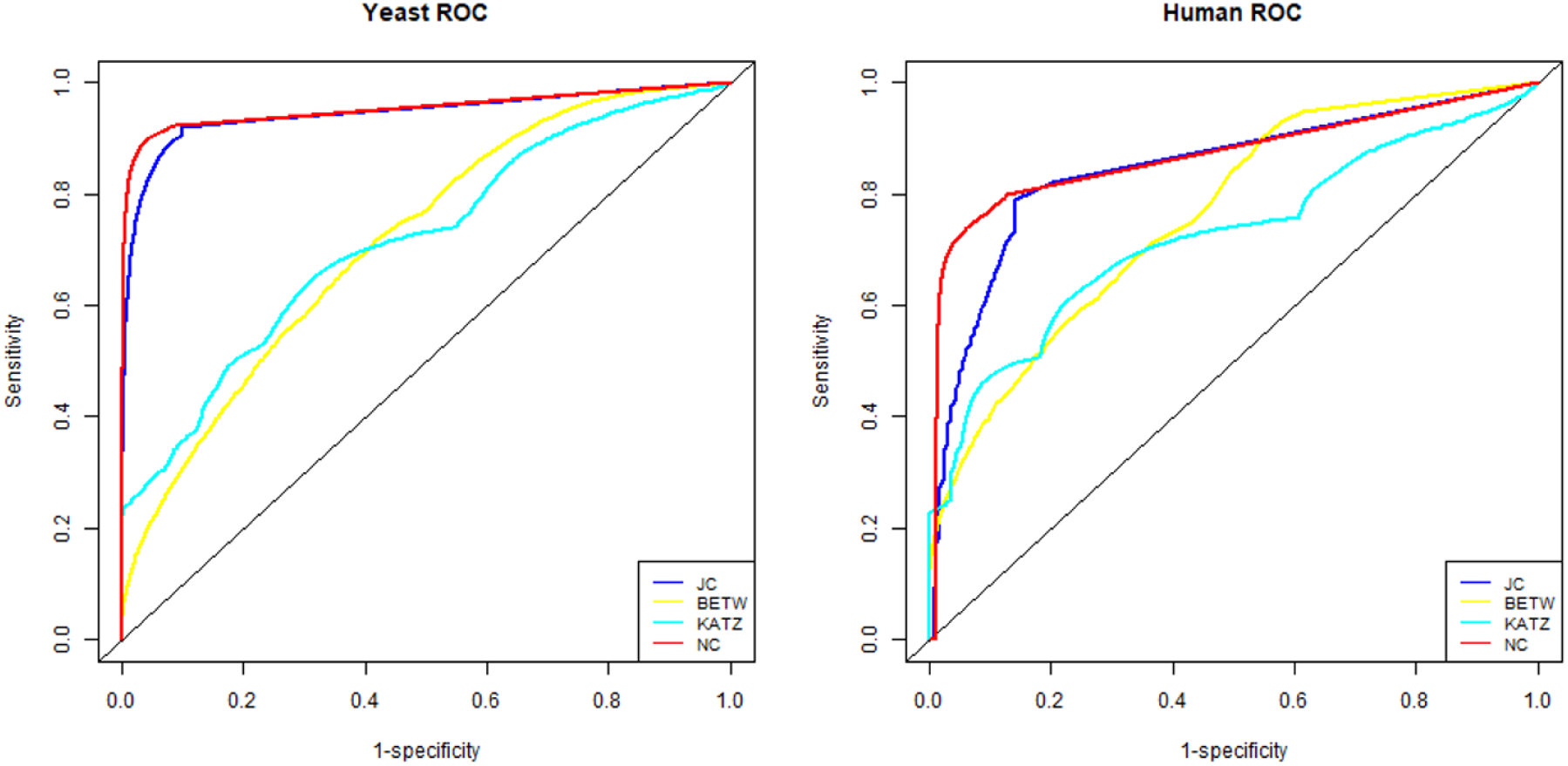
OM methodology ROC curves. ROC curves obtained by OM application with JC, BETW, KATZ and NC measures in Yeast and Human datasets.

**Figure 3.**
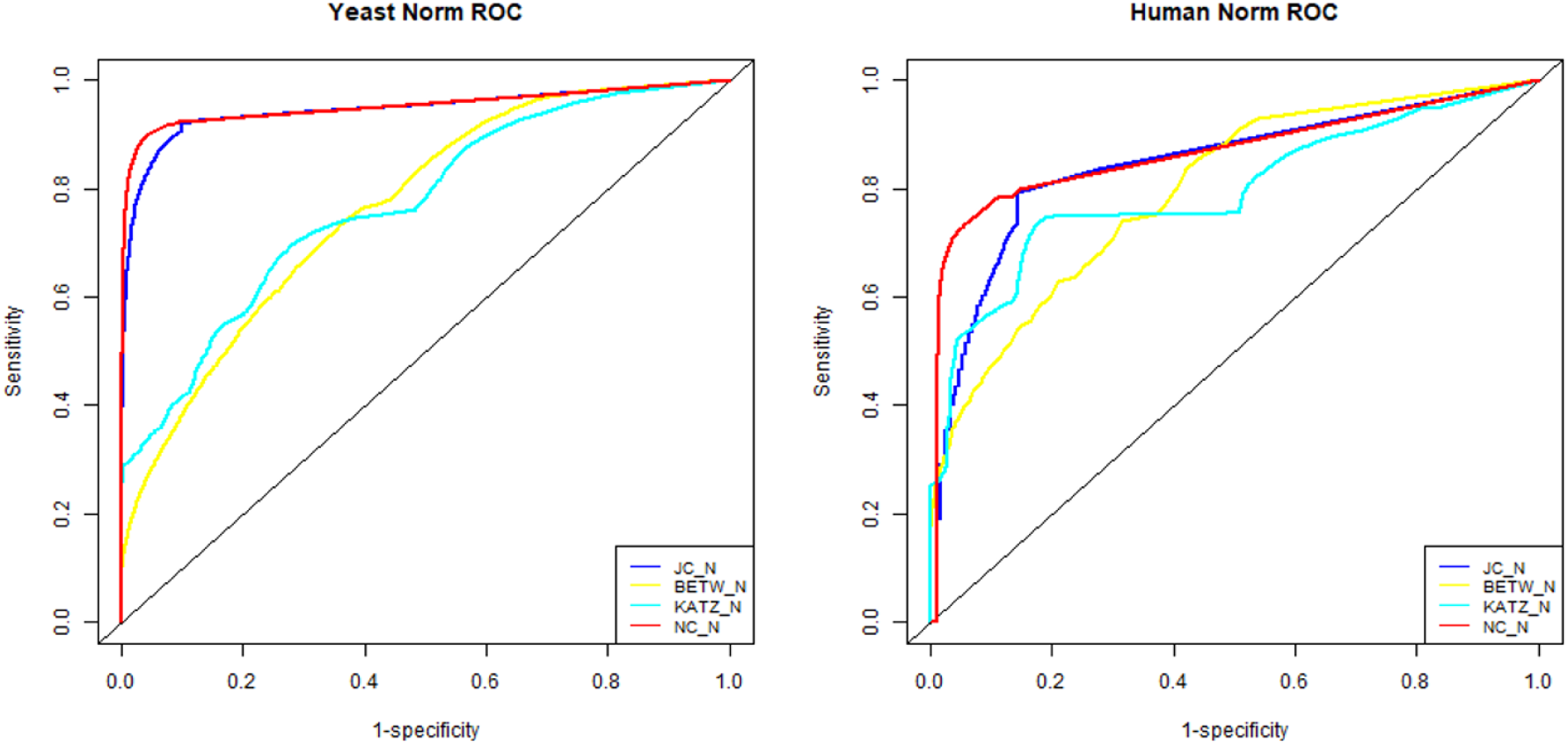
OM methodology ROC curves with normalized data. ROC curves obtained by OM application with JC, BETW, KATZ and NC measures in Yeast and Human datasets after data normalization between 0 and 1.

In addition to testing the OM methodology with these measures, we calculated the cut-off values for both the optimal cut and the accuracy cut, using the JC and NC measures, those that previously gave better results. The optimal cut calculates the point closest to (0,1) in the ROC curve, where sensitivity equals specificity, whereas the accuracy cut calculates the maximum accuracy and the respective cut-off value. In the next experiments, to determine OM THR, we calculated the cut-off values for the optimal cut, obtaining good results.

Table 3 and 4 show the obtained results for the Yeast and Human organisms. We can observe that the NC measure gave a slightly better result than JC and so we will describe forward the experiments using the NC measure.

**Table 3:** AUC, optimal cut and accuracy cut values in Yeast.

| Table 3: AUC, optimal cut and accuracy cut values in Yeast. |  |  |  |  |  |
| --- | --- | --- | --- | --- | --- |
| Yeast | AUC | Optimal cut |  | Accuracy cut |  |
| JC | 0.9462 | sensitivity | 0.9246 | accuracy | 0.9123 |
|  |  | specificity | 0.9000 |  |  |
|  |  | cut-off | 0.0008 | cut-off | 0.0008 |
| NC | 0.9534 | sensitivity | 0.9057 | accuracy | 0.9282 |
|  |  | specificity | 0.9471 |  |  |
|  |  | cut-off | 0.0021 | cut-off | 0.0044 |

**Table 4:**
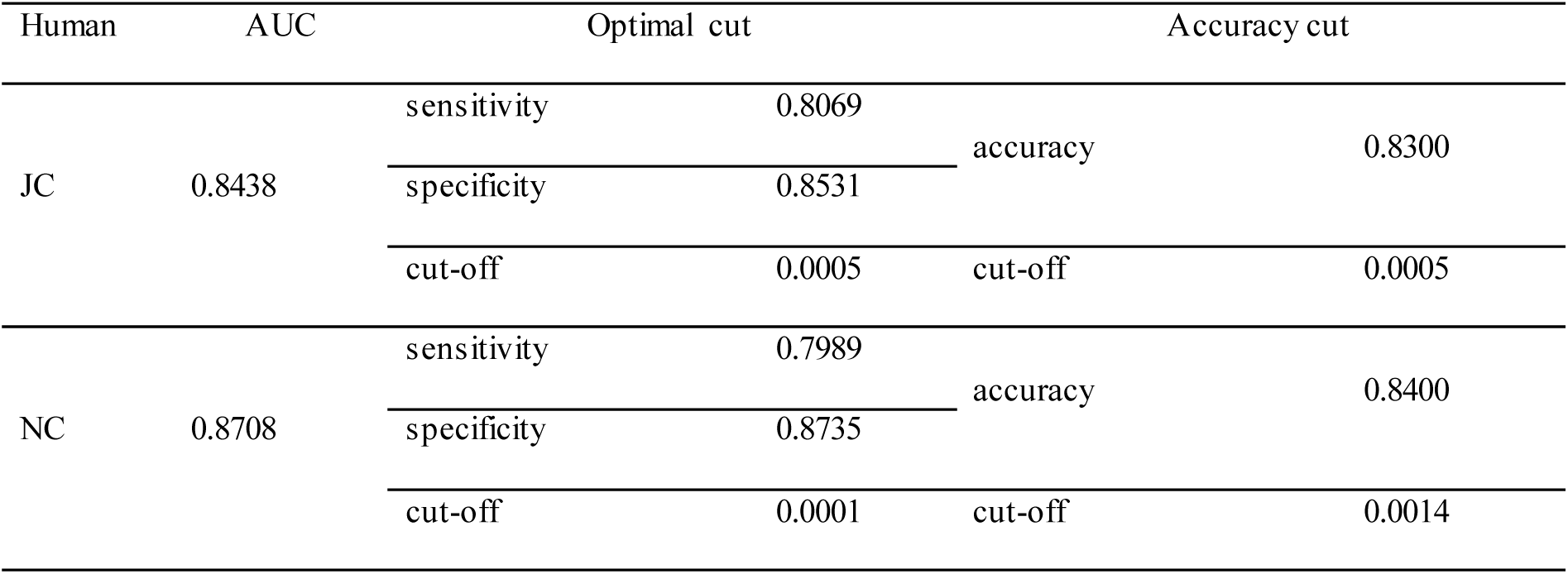
AUC, optimal cut and accuracy cut values in Human.

| Table 4: AUC, optimal cut and accuracy cut values in Human. |  |  |  |  |  |
| --- | --- | --- | --- | --- | --- |
| Human | AUC | Optimal cut |  | Accuracy cut |  |
| JC | 0.8438 | sensitivity | 0.8069 | accuracy | 0.8300 |
|  |  | specificity | 0.8531 |  |  |
|  |  | cut-off | 0.0005 | cut-off | 0.0005 |
| NC | 0.8708 | sensitivity | 0.7989 | accuracy | 0.8400 |
|  |  | specificity | 0.8735 |  |  |
|  |  | cut-off | 0.0001 | cut-off | 0.0014 |

### Evaluation of the organization measurement methodology in different scenarios

To assess whether the OM methodology is sensitive to the network topology, we applied it to a randomly generated protein network, with the same number of nodes and edges as their respective reference sets (for Yeast and Human). If the OM methodology can distinguish between interactions and non-interactions in the reference data sets but fails to do so in the random networks, one can assume that it captures the inherent topological structure of a real network.

To further evaluate the performance of OM methodology two other experiments were performed. First, while maintaining the same number of nodes (proteins), we randomly added incrementing percentages of edges (proteins interactions), 20%, 40%, 60% and 80%, not belonging to the reference set network, building four networks and removed the same percentages of edges from the reference set network, building more four networks. This was performed for the Yeast, Human and CS2007 reference set networks. After each addition or removal, we used the OM methodology to denoise the networks. To further assess the ability of the proposed methodology for network denoising, we also determined the percentage of inserted TN removed from the respective CS2007 perturbed networks and the percentage of TP retrieved from the respective CS2007 perturbed networks. A thorough description of the results of these experiments are shown in the following sub-sections.

### Organization measurement methodology performance comparison: random network versus reference set network

The only criteria selected to generate the random networks was that the resulting randomize networks were required to comprise the same number of nodes and edges. Thus, we generated 10 networks for each organism to be tested using the NC measure.

Fig 4 shows the ROC curves, the separation of classes (PPI e non-PPI) curves and the accuracy curve, when applying the OM methodology with the NC topological measure to one of the Yeast (left column) and Human (right column) random networks generated with the same number of nodes and edges of the respective reference sets. Analysing their ROC curves, we can see a clear distinction in performance between the application of OM methodology to the random network (Fig 4) and the subsets of the real networks (Fig 2). The AUC obtained after using the proposed method in all 10 random networks generated was close to 0.5 for both organisms (Yeast and Human), while for the subset of the Yeast network and Human network the AUC was 0.9534 and 0.8708, respectively.

**Figure 4:**
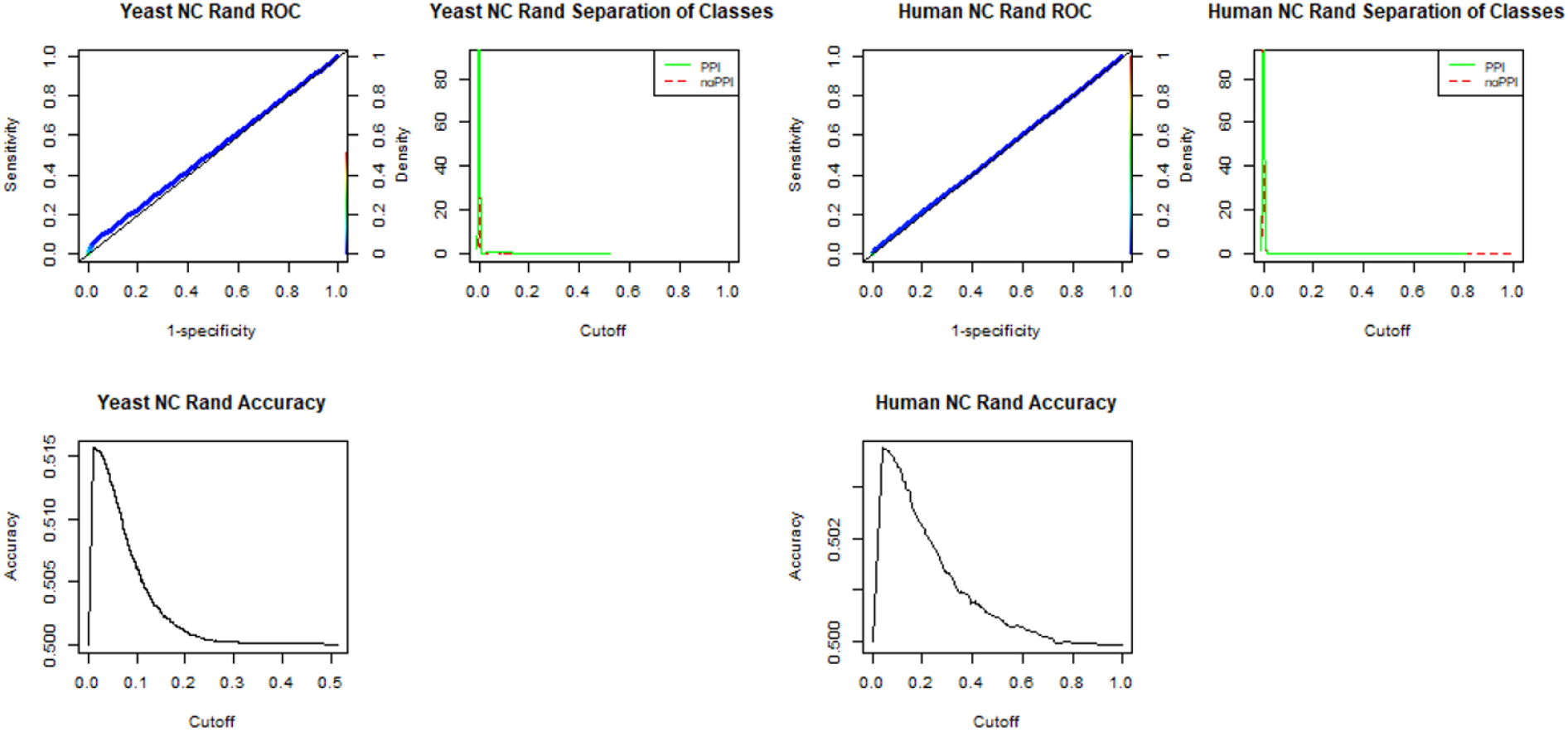
OM methodology application with the NC topological measure, to Yeast and Human random networks. OM methodology application ROC curves, separation of classes (PPI e non-PPI) curves and accuracy curve with the NC topological measure in one of the random networks generated with the same number of nodes and edges as the Yeast (left column) and as the Human (right column) reference sets.

### Random insertion of edges

To evaluate the performance of the OM methodology for denoising PPI networks, we perturbed the networks of the reference sets by randomly adding incrementing percentages of edges to the networks of the Yeast and Human reference sets and the CS2007 reference set.

We created four noisy networks for each data set, adding 20% more edges to the original network, followed by 40%, 60% and 80%. These intervals were selected following the research conducted by Yi Fang *et al.* [23]. Fig 5 shows the ROC curves when the OM methodology is applied to the networks of the Yeast and Human reference sets and to the four noisy networks generated from each of them. We can observe a decreasing of performance when we increase the percentage of the random edges added.

**Figure 5:**
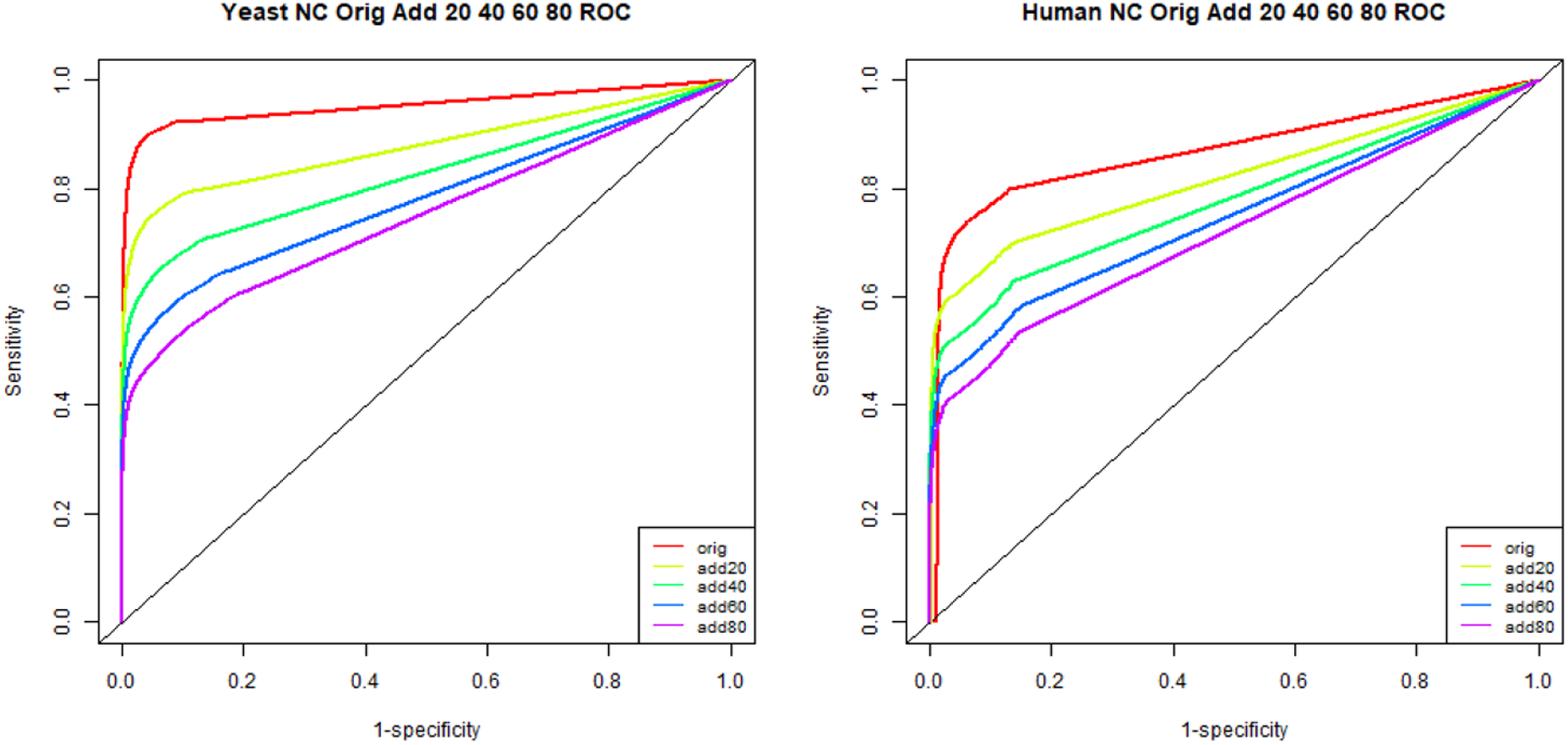
Application of the OM methodology with the NC topological measure, when an increasing percentage of edges was added randomly to the Yeast and Human reference set networks. ROC curves of the reference sets, and the other 4 networks, when 20%, 40%, 60% and 80% of edges were added to the reference set network for Yeast (left column) and Human (right column).

To be able to compare the performance of OM methodology with other network-based methodologies proposed by other researchers, we also perturbed the CS2007 network by randomly inserting edges in the same proportions previously described. Fig 6 shows the graphical representation of the resulting AUC values when the OM methodology is applied to the four noisy networks obtained from the CS2007 network and when MDS and IGS methodologies are applied [23]. It can be observed that the proposed OM methodology outperforms MDS and IGS methodologies.

**Figure 6:**
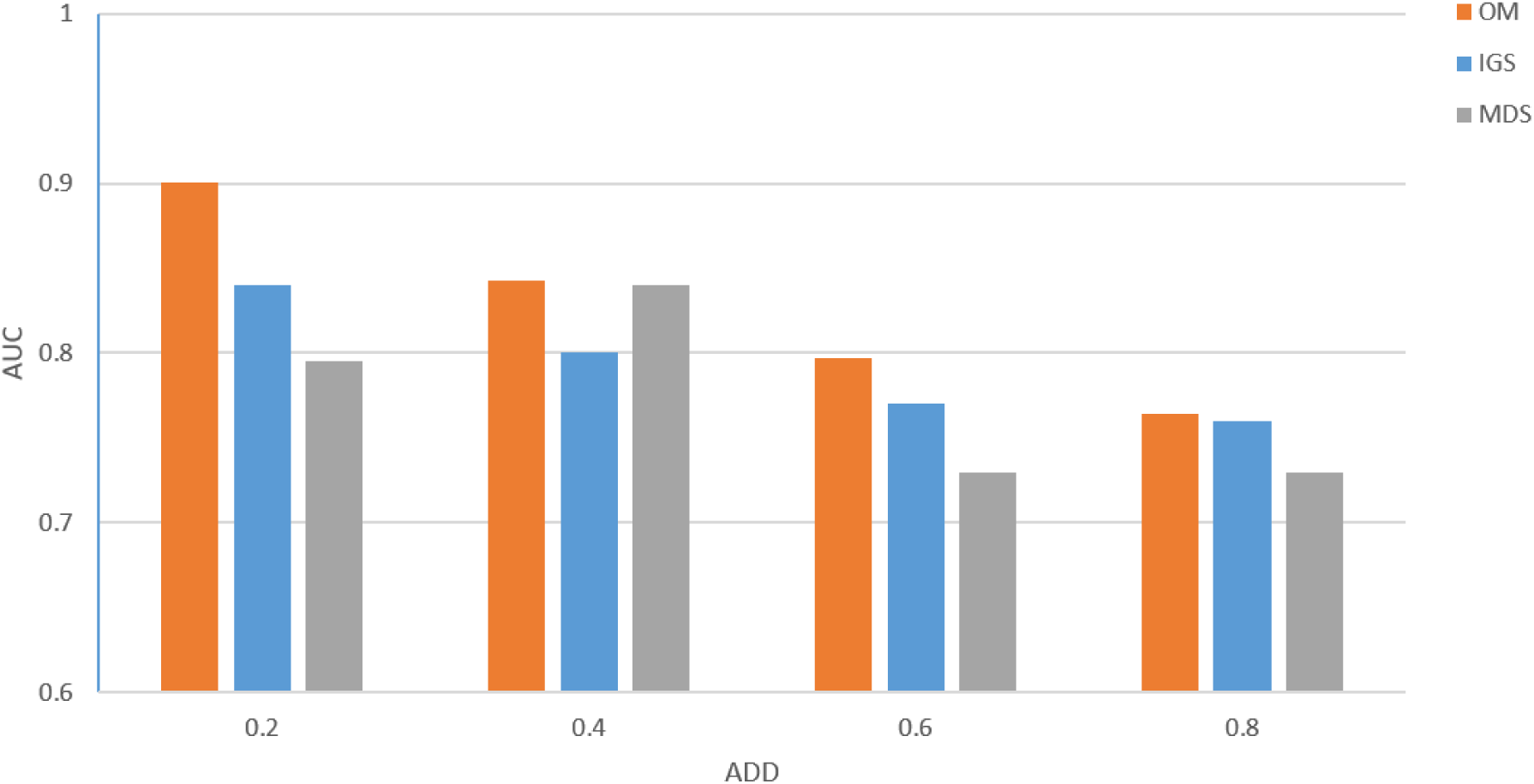
Application of the OM methodology with the NC topological measure, when an increasing percentage of edges was added randomly to the CS2007 reference set network compared to the MDS and IGS methods. AUCs values of the CS2007 perturbed 4 networks when a percentage of random 20%, 40%, 60% and 80% of edges were added to the reference set networks, using OM, IGS and MDS methods.

After denoising the networks with the OM methodology, we also calculated the percentage of FP interactions that were removed (Table 5). We observe that the OM methodology could remove 97% of the FP of the 20% added and 89% of the 80% added.

**Table 5:** Percentage of FP removed after applying the OM methodology to the noisy networks of the CS2007 dataset.

| % added | # FP added | # FP removed | % FP removed |
| --- | --- | --- | --- |
| 20 | 1,665 | 1,607 | 97 |
| 40 | 3,329 | 3,114 | 94 |
| 60 | 4,994 | 4,560 | 91 |
| 80 | 6,658 | 5,926 | 89 |

### Random deletion of edges

To evaluate the performance of the OM methodology for the identification of missing interactions, four new networks were created for each dataset (*i.e.*, Yeast, Human, and CS2007 reference sets) by removing increasing percentages of edges from the respective reference set networks. Edge removal was performed in the same proportion as edge addition: 20%, 40%, 60% and 80%. By removing increasing percentages of edges from the respective reference set networks, we are creating smaller sparse networks and at the same time we are deteriorating their inherent structure. The results are shown in Fig 7, representing the ROC curves, when the OM methodology is applied to the eight noisy networks referred previously, of the Yeast and Human reference set networks.

**Figure 7:**
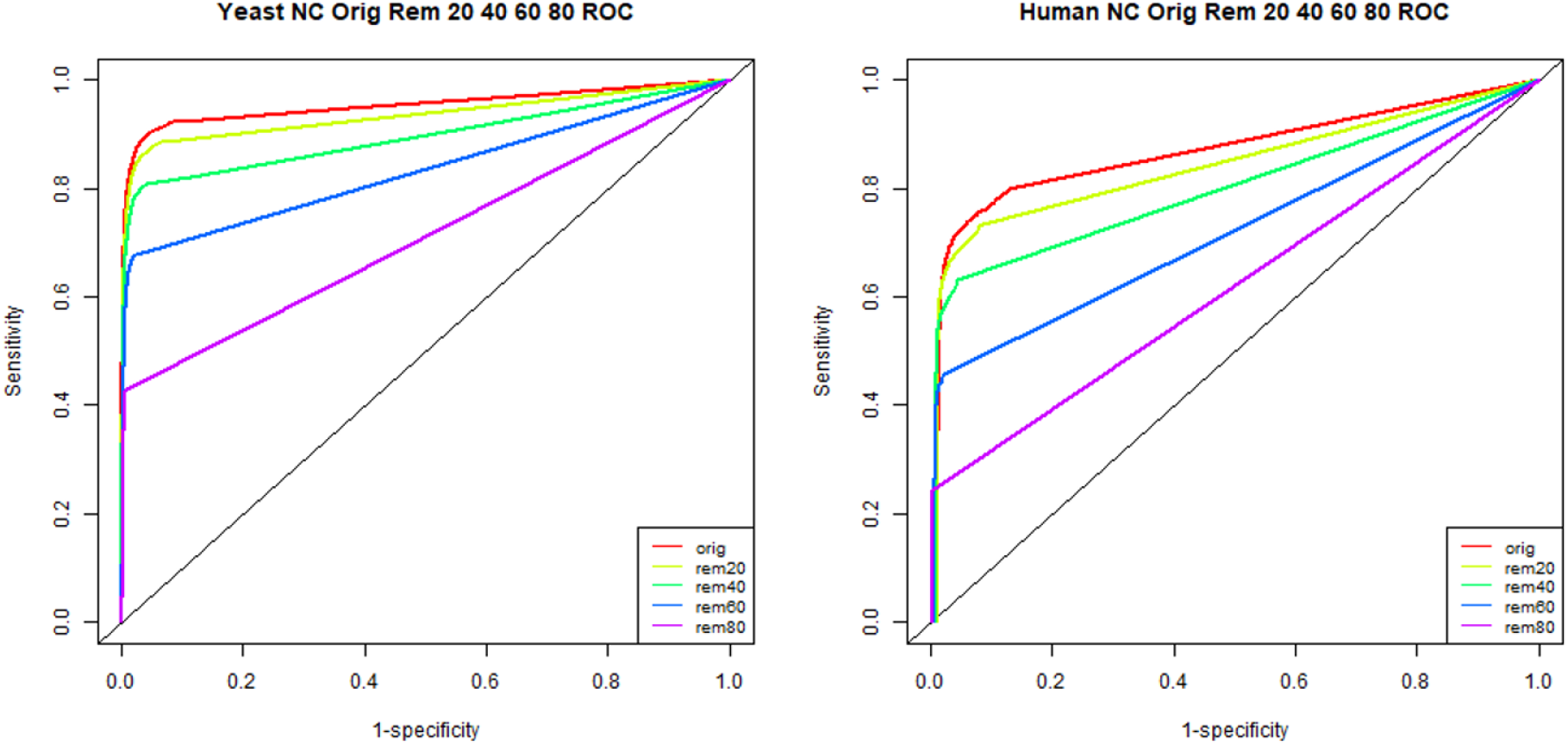
Application of the OM methodology with the NC topological measure, when an increasing percentage of edges is removed randomly from the Yeast and Human reference set networks. ROC curves of the reference set, and the other 8 networks when 20%, 40%, 60% and 80% of edges were removed from the reference set networks for Yeast (left column) and Human (right column).

These results show a scenario alike the one observed after randomly adding edges, as greater reductions in the number of edges result in greater performance drops, but the performance drops are steeper in the Human organism.

Fig 8 shows a graphic of the AUCs values, when the OM methodology is applied to the four noisy networks obtained from the CS2007 network, referred previously, compared to the MDS and IGS methodologies [23]. OM methodology has a better performance compared to the IGS and MDS methodologies, except when 80% of the interactions are removed from the CS2007 reference set, where the application of IGS gives better results. Further details are in Table 6, where we can observe that 95% of the TP removed could be detected when the OM methodology is applied to the perturbed network, when 20% of the interactions of the reference set were removed and 40% could be detected when 80% were removed.

**Table 6:** Percentage of TP inserted after applying the OM methodology to the incomplete networks of the CS2007 dataset.

| % removed | # TP removed | # TP inserted | % TP inserted |
| --- | --- | --- | --- |
| 20 | 1,665 | 1578 | 95 |
| 40 | 3,329 | 2993 | 90 |
| 60 | 4,994 | 4011 | 80 |
| 80 | 6,658 | 2633 | 40 |

**Figure 8:**
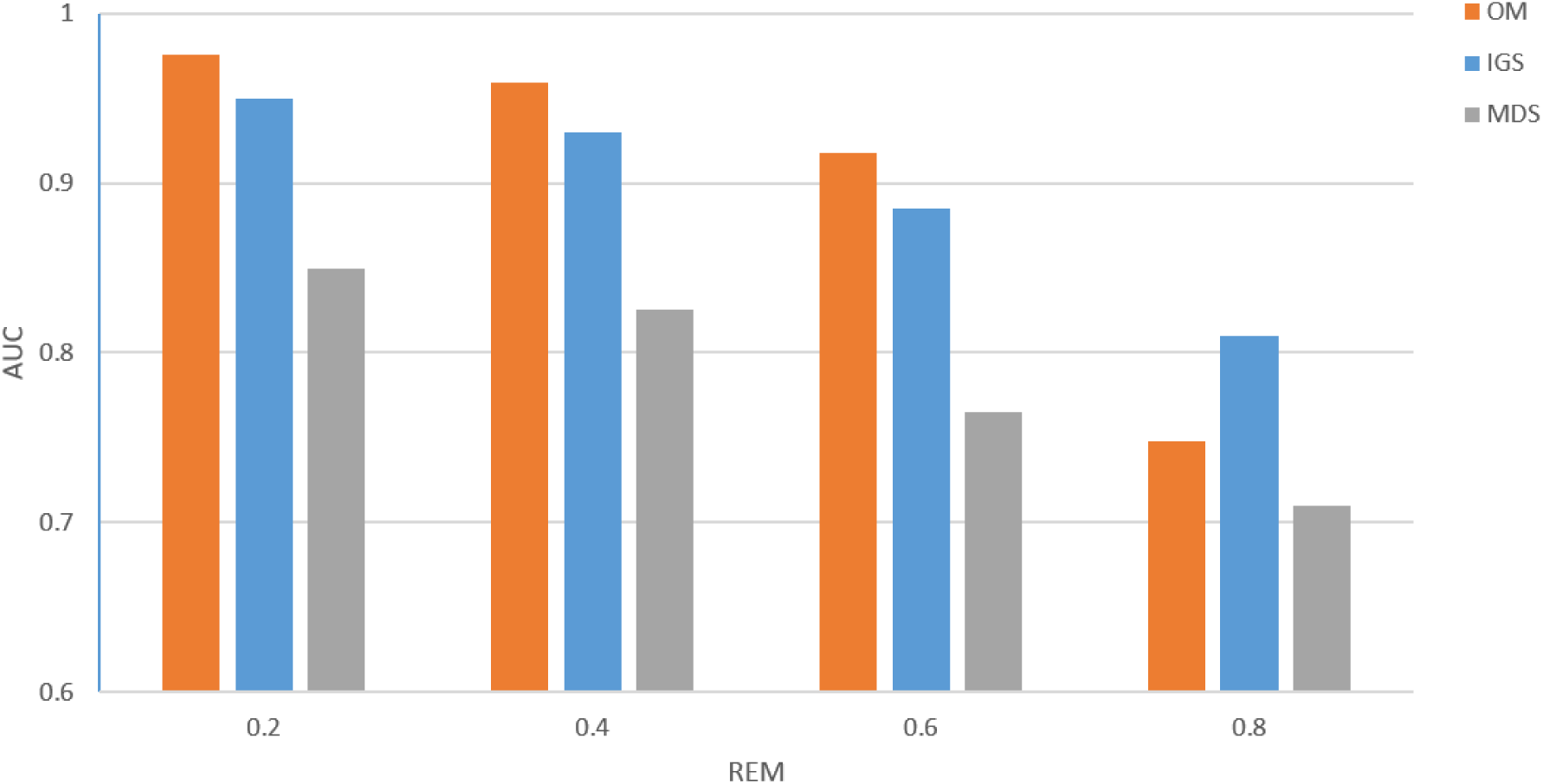
Application of the OM methodology with the NC topological measure, when an increasing percentage of edges was removed randomly to the CS2007 reference set network. AUCs values of the CS2007 perturbed 4 networks when 20%, 40%, 60% and 80% of edges were removed to the reference set networks, using OM, IGD and MDS methods.

### Analysis

Different topological measures were used to identify the optimal threshold, with the Yeast and Human reference sets and comparative testing showed (Figs 2 and 3 and Table 2) that the best results were obtained using the JC and the NC measures, and thus we decided to use both in some experiments of this work. JC is a widely known measure frequently used in network denoising and missing link prediction. It also considers the neighbourhood information, which is aligned with the “guilt-by-association” principle. The same applies to the NC index, proposed herein, where the concept of CC is also taken into account.

The OM methodology was then applied to the Yeast and Human datasets, using the JC and the NC measures and after analysing Tables 3 and 4, where the AUC values and the cut-off values for both the optimal cut and the accuracy cut for the Yeast and Human reference sets obtained are shown, it can be seen that the NC measure performed better than the JC measure at discriminating between protein interactions and non-interactions and for this reason the NC measure was used in the evaluation of the OM methodology.

Three different scenarios were considered to evaluate the OM methodology. The first one uses randomly generated protein networks, with the same number of nodes and edges as their respective reference sets (Yeast and Human). Observing the Fig 4 we can see that the AUC, obtained when applying the OM methodology to one of the random networks, was close to 0.5 for both organisms (Yeast and Human), while for the Yeast and Human reference sets, the AUC was 0.9534 and 0.8708, respectively (Fig 2), which shows that OM is sensitive to the inherent topological structure of a real network. These results show that the OM methodology cannot distinguish between interactions and non-interactions in random networks, but can capture the inherent rules of biological networks, not present in random networks.

The second scenario used to evaluate the performance of OM methodology, consisted in applying OM to networks obtained from the two Yeast and Human reference sets, where the number of nodes (proteins) was maintained, but where a random percentage of edges (proteins interactions), 20%, 40%, 60% and 80%, not belonging to the reference set network, were added, and the third scenario is similar to the second but instead of adding, the same random percentages of edges were removed from the reference set network.

In the second scenario (random insertion of edges), as expected, greater increments of random edges resulted in greater performance reductions (Fig 5). The performance reductions were steeper in Human, which could be attributed to one major reason: the percentage of FN is most likely greater in the Human interactome than in the Yeast interactome. Thus, it could be argued that the Yeast reference set is a more reliable, better representation of the actual Yeast interactome, than the Human reference set is of the real Human interactome. When we add these percentages of random edges, the inherent structure of these biological networks becomes deteriorated, because we are probably adding TN.

In the third scenario (random deletion of edges), greater reductions in the number of edges result in greater performance drops compared to the second scenario, but the performance drops are steeper in the Human organism (Fig 7). This could be explained by the fact that we are removing TP from both networks. However, since the Yeast network seems to be a closer representation of its true network than the Human network, the accentuated deterioration in the structure of the Human network could explain this behaviour.

So, when comparing the results between edge addition and edge removal in Yeast and Human reference sets (Figs 5 and 7), we witness that the overall performance reductions were quite dissimilar. Adding just 20% more edges contributed to a reduction of approximately 0.08 in AUC for Yeast, and 0.06 AUC for Human. Further addition of edges beyond this point did not decrease the AUC as sharply. Contrariwise, after removing 20% of the existing edges, the AUC decreased by roughly 0.02 for both Yeast and Human, with greater performance drops after each percentage of edges removal.

The better performance observed for the Yeast interactome could be explained by its smaller size compared to the Human interactome, in addition to being relatively well-studied, meaning that input data quality plays an important role in the performance of computational methods. Additionally, the negative impact in performance observed after randomly adding edges suggests that the OM methodology is very sensitive to high percentages of FP and FN.

To compare the performance of OM methodology with other network-based methodologies proposed by other researchers, CS2007 network reference set was perturbed by randomly inserting edges in the same proportions previously described in scenario two and by randomly deleting edges in the same proportions previously described in scenario three. OM methodology was compared to the MDS and IGS methodologies [23]. Figs 6 and 8 show the AUC values when these methodologies were applied to this dataset and it can be seen a general improvement in the performance when the OM methodology is applied compared to the MDS and IGS methodologies. When 80% of random TP of network interactions are removed, the AUC of IGS is superior than the AUC od OM and MDS. This could not mean, in this case, that OM is worst, since removing 80% of the TP makes the model very close to a random network and a denoising method, for consistency, should not be able to detect structure in such networks.

Further analysis was conducted for these last networks with added random percentages of FP interactions. OM methodology was applied and the percentage of FP interactions removed was calculated (Table 5). Interestingly, most of the randomly inserted FP interactions were promptly identified, even when the network was heavily perturbed, with 89% of the FP removed after contaminating the network with 6,658 random interactions. These results suggest that the OM methodology can indeed capture the inherent topology of biological networks. Interestingly, we observed that the number of TP identified after randomly removing edges from the CS2007 dataset plummets after removing 60% of TP (Table 6). Still, the OM methodology seems to identify most missing links up to that point. These findings suggest that the OM methodology can assess whether the topological structure of a network is according to the characteristic topology of biological networks.

OM methodology could still work well in less-studied interactomes, when the subset of the interactome of interest is a representative sample of the structure of the entire interactome, meaning that the percentage of FP and FN cannot hide the inherent structure behind the biological networks of the organisms.

## Conclusions

Currently, low-throughput experimental methods are the only effective way to validate protein interactions. While high-throughput experimental methods to obtain PPIs exist, the obtained results have very high noise. As such, computational methods are required to speed data acquisition and to reduce the data contamination. Methods relying exclusively in the topology of biological networks are simpler and faster, as it appears that networks topology may reveal patterns or signatures associated with the kind of organism and the type of interactions. If we can use, effectively, only the topology to denoise biological networks, we have a simple computational method suitable for incomplete interactomes, without the need of extra biological knowledge.

This paper introduces the OM methodology for denoising biological networks, a methodology that: a) uses exclusively the topology of the network; b) enables, easily, to separate the distributions of interaction and non-interaction proteins in PPI networks; c) does not use known distributions as approximations; and d) provides a topological way of detecting FP interactions and find new interactions. The main innovation of the OM methodology is related with its ability to combine the advantages of using exclusively the topology without taking approximations to known distributions and without using external knowledge to detect interactions that do not exist or to find new interactions with a better performance than some documented used methodologies. This paper also introduces a new network topologic measure, the NC measure, which is used with the OM methodology and yielded better results, compared to other known and current topological measures.

The OM methodology can be explored in the future by applying it in networks belonging to other domains, where there is an inherent structure, to predict new interactions and eliminate spurious interactions.

(Preprint: doi: https://doi.org/10.1101/527606)

## Data Availability

The datasets supporting the conclusions of this article are available in the Bioinformatics repository of the University of Aveiro, at http://bioinformatics.ua.pt/software/OM.

## Acknowledgements

Not applicable.

